# Imputing not available values in single-cell DNA methylation data using the median is straightforward and effective

**DOI:** 10.1101/2024.10.17.618821

**Authors:** Songming Tang, Siyu Li, Shengquan Chen

## Abstract

Recent advances in single-cell DNA methylation have provided unprecedented opportunities to explore cellular epigenetic differences with maximal resolution. Due to the number of methylation sites exceeding the computational limits of current analytical methods, a common workflow for single-cell DNA methylation analysis is binning the genome into multiple regions and computing the average methylation level within each region. In this process, imputing not available (NA) values which are caused by the limited number of captured methylation sites is a necessary preprocessing step for downstream analyses. Existing studies have employed several simple imputation methods (such as zero imputation or mean imputation); however, there is a lack of theoretical studies or benchmark tests evaluating these approaches. Through both experiments and theoretical analysis, we found that using the median to impute missing data can effectively and simply reflect the methylation state of the NA values, providing an accurate foundation for downstream analyses.

---

DNA methylation (DNAm) is one of the earliest identified types of epigenetic modification and plays an essential role in regulating normal cellular processes, embryogenesis, and tumor development and progression^1, 2^. Recent advances in single-cell DNA methylation (scDNAm) have provided unprecedented opportunities to explore cellular epigenetic differences with maximal resolution. Most current studies analyze single-cell DNA methylation data typically based on cell-by-region matrix^3-5^. One simple yet effective method for scDNAm data creating cell-by-region matrices is genome window binning, which aggregates signals and simplifies the analysis^3, 6-8^. By binning the genome into tiles of fixed lengths, such as 100 kbp, and computing the average DNA methylation level for each cell in each region, a cell-by-region methylation matrix can be constructed.

Before conducting downstream analyses, a critical issue must still be addressed: handling the not available values inherent in scDNAm data. For single-cell RNA sequencing (scRNA-seq) or single-cell assay for transposase-accessible chromatin using sequencing (scATAC-seq), dropouts in sequencing lead to read counts of zeros^5^. However, in scDNAm data, captured methylation sites typically display a binary characteristic: methylated (read count of 1) or unmethylated (read count of 0), while uncaptured sites are not available (Fig. 1A). When constructing a cell-by-region matrix using the window binning strategy, due to the uneven distribution of methylation sites across the genome and the effect of window size, a considerable number of regions will still have no captured methylation sites and have average methylation level marked as not available (NA) values (referred to as missing values). A methylation matrix with NA values is not permitted for downstream analysis, making the imputation of the methylation matrix a necessary preprocessing step (Fig. 1A).

**Fig. 1.**
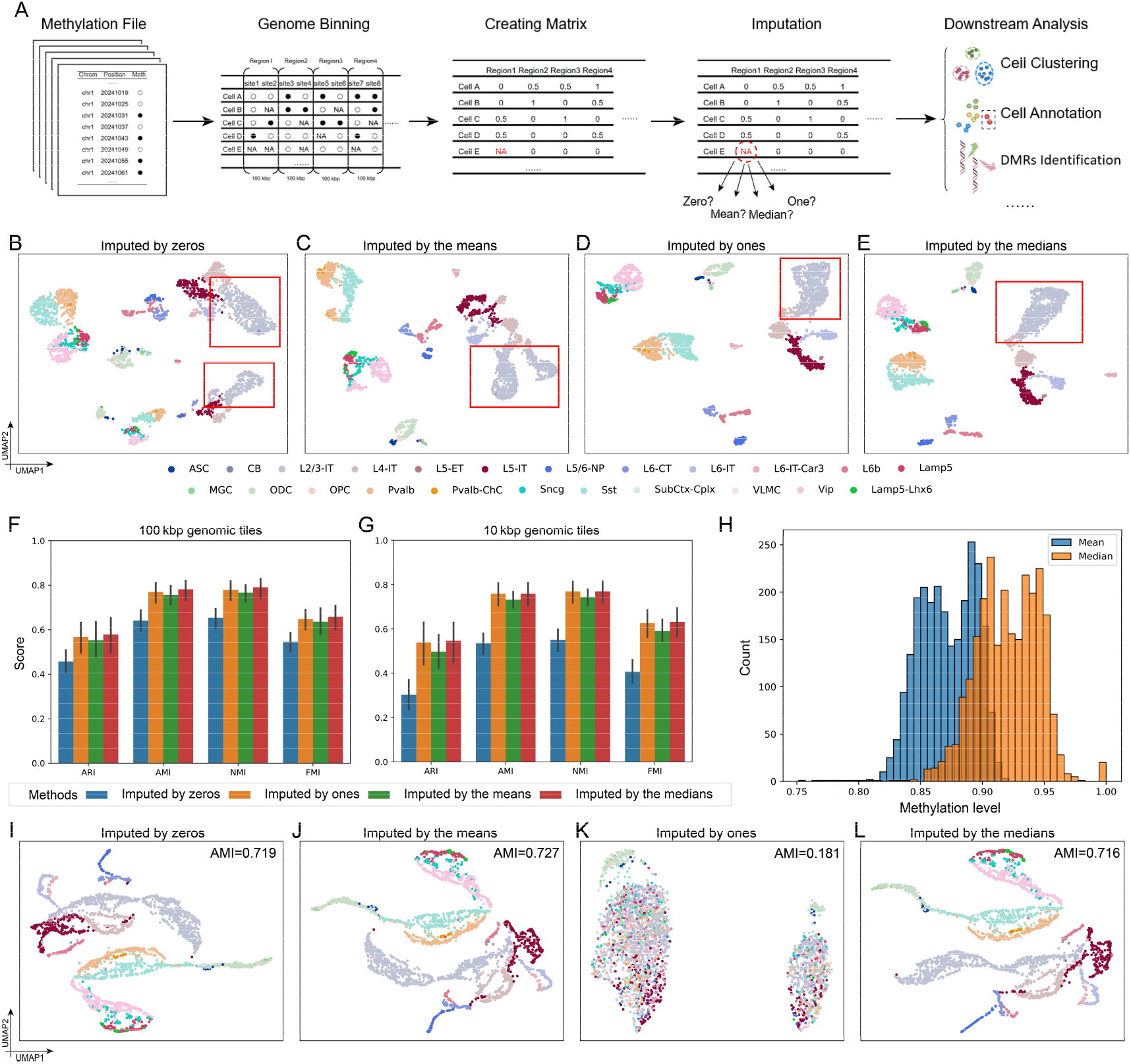
The effect of various imputation strategies. **A**, Workflow of single-cell DNA methylation data preprocessing. **B-E**, UMAP visualizations of the CG methylation on “GSE167577” dataset with a region length of 100 kbp imputed by **B**, zeros, **C**, the means, **D**, the ones and **E**, medians. **F**, Clustering performance across 11 CG methylation datasets, evaluated using various metrics for different imputation strategies at tiles of 100 kbp. **G**, Clustering performance across 11 CG methylation datasets, evaluated using various metrics for different imputation strategies at tiles of 10 kbp. **H**, Boxplot illustrating the distribution of the means and the medians of the “GSE167577” dataset at a region length of 100 kbp. **I-L**, UMAP visualizations and clustering performance of the CH methylation on “GSE167577” dataset with a region length of 100 kbp imputed by **I**, zeros, **J**, the means, **K**, the ones and **L**, the medians. The legend of **I-L** is the same as **B-E**.

When analyzing scDNAm data in a manner analogous to scRNA-seq data and scATAC-seq data, an intuitive solution would be imputing all NA values as zeros. For example, Luo et al.^6^ and Acharya et al.^9^ impute regions without any DNA methylation signal as zeros in their data processing pipelines. However, from an alternative perspective, in scRNA-seq data, higher read counts correspond to higher gene expression levels, while gene expression is strongly negatively correlated with DNA methylation levels^8^. The dropouts in scRNA-seq data often indicate a high methylation level; therefore, the dropout values in scRNA-seq data are treated as zero, which is equivalent to imputing the NA values in scDNAm data as ones. Additionally, utilizing various statistical measures to smooth NA values presents an intuitive approach. Episcanpy imputes NA values using the means of methylation levels in a region across all cells, thus avoiding NA issues^4^. Although these studies provide several imputation strategies, in our research, we found that imputing with the median can more effectively impute the NA values in scDNAm data and improve downstream analysis workflows.

We conducted comprehensive tests on 11 datasets with different sources, protocols, and species to evaluate the effect of various imputation strategies of scDNAm data^3, 8^ (Supplementary Table S1). Following the DNA methylation data analysis workflow of Episcanpy, we created methylation matrices for each dataset using window binning of a fixed length at the whole-genome level^4^. Subsequently, we applied four different imputation strategies to impute the NA values: using zeros, ones, the means, and the medians (referring to the means or medians methylation level across all cells in a given region). We then performed dimensionality reduction using principal component analysis (PCA) to reduce the data to 50 dimensions and applied the Louvain algorithm with default parameters for unsupervised cell clustering. An effective imputation strategy should better characterize cellular heterogeneity and improve clustering as well as other downstream analyses. Since the datasets were properly annotated, we adopted widely used metrics including the adjusted Rand index (ARI), adjusted mutual information (AMI), normalized mutual information (NMI), and Fowlkes-Mallows index (FMI) to evaluate clustering performance (Supplementary Note S1).

Methylation of vertebrate genomes mainly occurs on CG dinucleotides, research on CG methylation is a primary focus in current studies^1^. Thus, we first investigated the performance of different imputation strategies on CG methylation data. Following Tian et al., we first generated cell-by-region matrices with a region length of 100 kbp^8^. We used uniform manifold approximation and projection (UMAP) to visualize the embeddings obtained by PCA from the data imputed by different methods. Taking the “GSE167577” dataset as an example, the visualization of the data imputed by zeros or the means showed that many cell types, such as L4-IT, L5-IT, and L6-IT, were incorrectly split into two distinct clusters (Fig. 1B,C). Moreover, data imputed by the medians or ones successfully distinguished the same cell types together and accurately captured the heterogeneity among cells from different cell types (Fig. 1D,E). The clustering results at 100 kbp tiles of CG methylation in 11 datasets also showed that imputation using the medians and ones yielded the best performance. Across various metrics, data imputed by the medians showed 2% advantage over the data imputed by ones (Fig. 1F). Moreover, imputation by the means performed slightly worse, with the medians outperforming the means by 4.56% in ARI and 3.27% in AMI (Fig. 1F). Imputation with zeros performed significantly worse than the other methods, with its clustering metrics showing a greater than 20% disadvantage compared to the medians (Fig. 1F).

To investigate imputation strategies at higher region resolutions, we reduced the region length to 10 kbp and repeated the above experiments. The results indicated that the medians and one imputation were almost equivalent and highly effective for handling NA values at shorter tiles. However, the disadvantages of mean imputation became more pronounced at the 10 kbp level, with a 10% reduction in ARI compared to median or one imputation (Fig. 1G). Compared to the 100 kbp tiles, the 10 kbp tiles introduced more severe data missing, with most available values deviating significantly from zero (Fig. 1G). As a result, using zeros to impute NA values introduced substantial erroneous signals, leading to a 44.61% drop in ARI and a 29.49% drop in AMI compared to median imputation (Fig. 1G). This indicates that zero imputation is highly unsuitable for shorter tiles because the zero imputation in increased missing data can introduce more noise.

It is worth noting that in most single-cell DNA CG methylation datasets, because of the high methylated proportion of CG sites, the majority of regions exhibit high methylation levels. In such cases, the median is often close to one (Fig. 1H). The mean tends to be lower than the median due to the presence of differentially methylated regions (DMRs) with lower methylation levels (Fig. 1H). This causes the mean to deviate from the average methylation level of NA values. Specifically, these lower methylation regions decrease the mean more significantly, whereas the median is less affected due to the relative rarity of these low-methylation DMRs.

For CH methylation data, we used a region length of 100 kbp and conducted experiments on “GSE167577” datasets. The results of imputing with the medians for CH methylation data consistently showed favorable performance (Fig. 1I-L). However, due to the extremely low methylated proportion of CH sites, using ones for imputation introduced significant noise, resulting in a complete loss of cellular heterogeneity in the imputed data (Fig. 1K).

Given the above considerations and experimental results, we conclude that although imputing NA values as zeros can still reveal cell-type specificity in scDNAm data to some extent, it is not the recommended approach for handling NA values in scDNAm data. Moreover, while imputing with means is a widely used and generally effective method, it is significantly influenced by DMRs, making it unable to accurately reflect the average methylated state of NA values. Furthermore, due to the high methylated rate of CG sites and the strong negative correlation between CG methylation level and gene expression, using ones for imputing CG methylation data is feasible. However, for CH methylation, which has an extremely low methylated rate, imputing with ones would introduce intolerable noise. In conclusion, imputing NA values with the medians best exhibits cellular heterogeneity and preserves the biological signal.

In summary, based on our findings, we suggest that imputing NA values with the median is a straightforward and effective way to highlight cellular heterogeneity in scDNAm data, offering an accurate data foundation for downstream analyses and allowing for a more accurate and reliable interpretation of the underlying biology.

## Compliance and ethics

The authors declare that they have no conflict of interest.

## Acknowledgments

This work was supported by the National Natural Science Foundation of China [62203236, 62473212], the Young Elite Scientists Sponsorship Program by CAST [2023QNRC001]

