## Supplementalary files for "Imputing not available values in single-cell DNA methylation data using the median is straightforward and effective"

**Supplementary Note S1: Details of the evaluation metrics.**

For ARI, suppose that $C$ are the actual labels of the clusters and $K$ are the clustering labels. In that case, we define $a$ and $b$ as follows: $a$, the number of pairs of elements in the same set in $C$ and the same set in $K$. $b$, the number of pairs of elements in different sets in $C$ and different sets in $K$. The following formula then gives the original Rand index:

$$\begin{aligned} \begin{matrix} RI=\frac{a+b}{C_{2}^{n_{\text{samples }}}} \end{matrix} \#\left( 1 \right) \end{aligned}$$

$C_{2}^{n_{\text{samples }}}$is the total number of possible pairs in the datasets, and the following formula gives the ARI, which ensures random matching will get a value close to 0 and perfect matching gets a value close to 1:

$$\begin{aligned} \begin{matrix} ARI=\frac{RI-E\left[ RI \right]}{\max\left( RI \right)-E\left[ RI \right]} \end{matrix} \#\left( 2 \right) \end{aligned}$$

Mutual Information (MI) evaluates the degree of dependence between two random variables. AMI adjusts MI by considering the expected value under random clustering, as shown in Equation $\left( 3 \right)$.

$$\begin{aligned} \begin{matrix} AMI=\frac{MI\left( \mathbf{P},\mathbf{T} \right)-E\left[ MI\left( \mathbf{P},\mathbf{T} \right) \right]}{avg\left[ H\left( \mathbf{P} \right),H\left( \mathbf{T} \right) \right]-E\left[ MI\left( \mathbf{P},\mathbf{T} \right) \right]} \end{matrix} \#\left( 3 \right) \end{aligned}$$

Besides, the Fowlkes-Mallows Index (FMI) can be used as the geometric mean of pairwise precision and recall.

$$\begin{aligned} FMI=\frac{TP}{\sqrt{\left( TP+FP \right)\left( TP+FN \right)}}\#\left( 4 \right) \end{aligned}$$

**Supplementary Table S1.** Summary of 11 CG methylation datasets used in this study.

| **Dataset** | **Species** | **Sequencing Method** | **No. of cells** | **No. of cell types** | **NA values**  **proportion (100 kbp)** | **NA values**  **proportion (10 kbp)** | **Literature** |
| --- | --- | --- | --- | --- | --- | --- | --- |
| GSE130553 | *Mus musculus* | snmC-seq2 | 1367 | 22 | 2.94% | 19.47% | Liu et al.^1^ |
| GSE131354 | *Mus musculus* | snmC-seq2 | 296 | 21 | 4.55% | 16.33% | Liu et al.^1^ |
| GSE131360 | *Mus musculus* | snmC-seq2 | 1262 | 22 | 5.73% | 18.88% | Liu et al.^1^ |
| GSE131393 | *Mus musculus* | snmC-seq2 | 1106 | 26 | 2.85% | 21.96% | Liu et al.^1^ |
| GSE131406 | *Mus musculus* | snmC-seq2 | 1433 | 24 | 2.84% | 26.50% | Liu et al.^1^ |
| GSE167577 | *Homo sapiens* | snmC-seq3 | 2518 | 23 | 2.24% | 33.37% | Tian et al.^2^ |
| GSE168066 | *Homo sapiens* | snmC-seq3 | 2496 | 21 | 2.23% | 31.83% | Tian et al.^2^ |
| GSE168645 | *Homo sapiens* | snmC-seq3 | 2951 | 22 | 2.39% | 35.96% | Tian et al.^2^ |
| GSE168734 | *Homo sapiens* | snmC-seq3 | 2496 | 21 | 2.23% | 30.64% | Tian et al.^2^ |
| GSE179610 | *Homo sapiens* | snmC-seq3 | 2786 | 20 | 1.69% | 22.53% | Tian et al.^2^ |
| GSE179971 | *Homo sapiens* | snmC-seq3 | 2730 | 25 | 2.29% | 34.89% | Tian et al.^2^ |
